# Genetically encoded ratiometric indicators for potassium ion

**DOI:** 10.1101/254383

**Authors:** Yi Shen, Sheng-Yi Wu, Vladimir Rancic, Yong Qian, Shin-Ichiro Miyashita, Klaus Ballanyi, Robert E. Campbell, Min Dong

## Abstract

Potassium ion (K^+^) homeostasis and dynamics play critical roles in regulating various biological activities, and the ability to monitor K^+^ spatial-temporal dynamics is critical to understanding these biological functions. Here we report the design and characterization of a Förster resonance energy transfer (FRET)-based genetically encoded K^+^ indicator, KIRIN1, constructed by inserting a bacterial cytosolic K^+^ binding protein (Kbp) between a fluorescent protein (FP) FRET pair, mCerulean3 and cp173Venus. Binding of K^+^ induces a conformational change in Kbp, resulting in an increase in FRET efficiency. KIRIN1 was able to detect K^+^ at physiologically relevant concentrations *in vitro* and is highly selective toward K^+^ over Na^+^. We further demonstrated that KIRIN1 allowed real-time imaging of pharmacologically induced depletion of cytosolic K^+^ in live cells, and KIRIN1 also enabled optical tracing of K^+^ efflux and reuptake in neurons upon glutamate stimulation in cultured primary neurons. These results demonstrate that KIRIN1 is a valuable tool to detect K^+^ *in vitro* and in live cells.

## Introduction

Förster resonance energy transfer (FRET)-based genetically encoded indicators are widely used tools for studying biochemical activities in live cells^1,2^. Intramolecular FRET indicators are particularly useful for the detection of binding-induced protein conformational change^3^. The design principle of such indicators is straightforward and well-established: a sensing domain is attached to two fluorescent proteins (FPs) in a single polypeptide form. Upon analyte binding, the conformational change of the sensing domain affects the FRET efficiency between the attached fluorophores, altering the ratiometric fluorescence emission^4^. The development of FP FRET-based genetically encoded indicators for Ca^2+^, Zn^2+^, Mg^2+^, ATP, neurotransmitter, membrane voltage, and enzyme activities has opened new avenues for biological studies and led to numerous new biological insights^5–13^. However, analogous indicators for monovalent metal cations including sodium (Na^+^) and potassium (K^+^) ions have been noticeably absent from this toolkit until very recently.

The measurement of K^+^ concentration in biological systems is critical, considering the large impact of K^+^ levels on all aspects of cellular homeostasis^14^. Normal levels of K^+^ concentration ( ∼150 mM for intracellular K^+^; ∼5 mM for extracellular K^+^) are vital for the maintenance of neuronal^15,16^, cardiovascular^17^, as well as immune system function^18–20^, while abnormal K^+^ concentration levels are often associated with certain disease conditions^21,22^. Measuring K^+^ concentrations has predominantly relied upon K^+^-specific glass capillary electrode^23^. Although sensitive and accurate, such electrode-based measurements are invasive for patients, time consuming, and low throughput. Electrode-based measurements also provide little to no spatiotemporal information on K^+^ dynamics in biological samples. Alternatively, synthetic small molecule-based K^+-^sensitive fluorescent dyes have been developed to measure K^+^ concentration, but these dyes usually have poor selectivity, as they also bind to Na^+24,25^. Although K^+^-sensitive dyes with improved selectivity have been reported^26,27^, the use of synthetic dyes still involves complications including cumbersome loading and washing steps. In addition, synthetic dyes cannot be targeted to specific cells in a tissue.

Developing genetically encoded K^+^ indicators will greatly facilitate the study of K^+^ homeostasis and dynamics in live cells and *in vivo,* and will allow accurate measurement of K^+^ concentration levels in specific cell types or cellular organelles with spatial and temporal resolution. The key to designing such an indicator is to identify or develop a suitable sensing domain with a high degree of specificity toward K^+^ and sufficient levels of conformational change upon binding to K^+^. Recently, an *E. coli* K^+^ binding protein (Kbp) was identified and structurally characterized^28^. Kbp is a small (149 residues, 16 kDa) protein with highly specific cytoplasmic K^+^ binding properties. Kbp contains two domains: BON (bacterial OsmY and nodulation)^29^ at the N-terminus and LysM (lysin motif)^30^ at the C-terminus. Importantly, structural analysis of Kbp has revealed that the protein exhibits a global conformational change upon K^+^ binding^28^, bringing BON and LysM closer to each other to stabilize K^+^. This high level of K^+^ specificity and large degree of conformational change are ideal properties for constructing a genetically encoded K^+^ indicator. Building upon this insight, we have designed and developed genetically encoded K^+^ indicators that utilize Kbp and FP-based FRET pairs.

## Results

### Design and construction

We designed genetically encoded K^+^ indicators by inserting Kbp between a FP FRET pair (Fig. 1a). We expected that, upon binding of K^+^, the Kbp sensing domain would undergo a conformational change, increasing FRET efficiency between the FP donor and acceptor (Fig. 1b). Accordingly, we would observe a ratiometric fluorescence change with a donor emission fluorescence decrease and an acceptor emission fluorescence increase. We constructed a fusion protein consisting of two FPs: the cyan FP (CFP) mCerulean3 as the FRET donor^31^; and a circularly permutated variant of yellow FP (YFP), cp173Venus^32^, as the FRET acceptor. We chose mCerulean3 because it is the brightest CFP currently available^33^ with an exceptionally high quantum yield at 0.80; and cp173Venus is a bright FRET acceptor with orientation factor optimized for a CFP-based donor^32^. mCerulean3 is linked to the N-terminus of Kbp (residues 2-149) with cp173Venus attached to the C-terminus (Fig. 1a, b). The resulting K^+^ indicator was designated KIRIN1 (potassium/K ion ratiometric indicator). We also construct a red-shifted K^+^ indicator with an alternative fluorescence hue, using a FRET pair of the green FP (GFP) Clover and the red FP (RFP) mRuby2^34^ (Supplementary Fig.1). The resulting indicator was named KIRIN1-GR (KIRIN1 with GFP and RFP).

**Figure 1.**
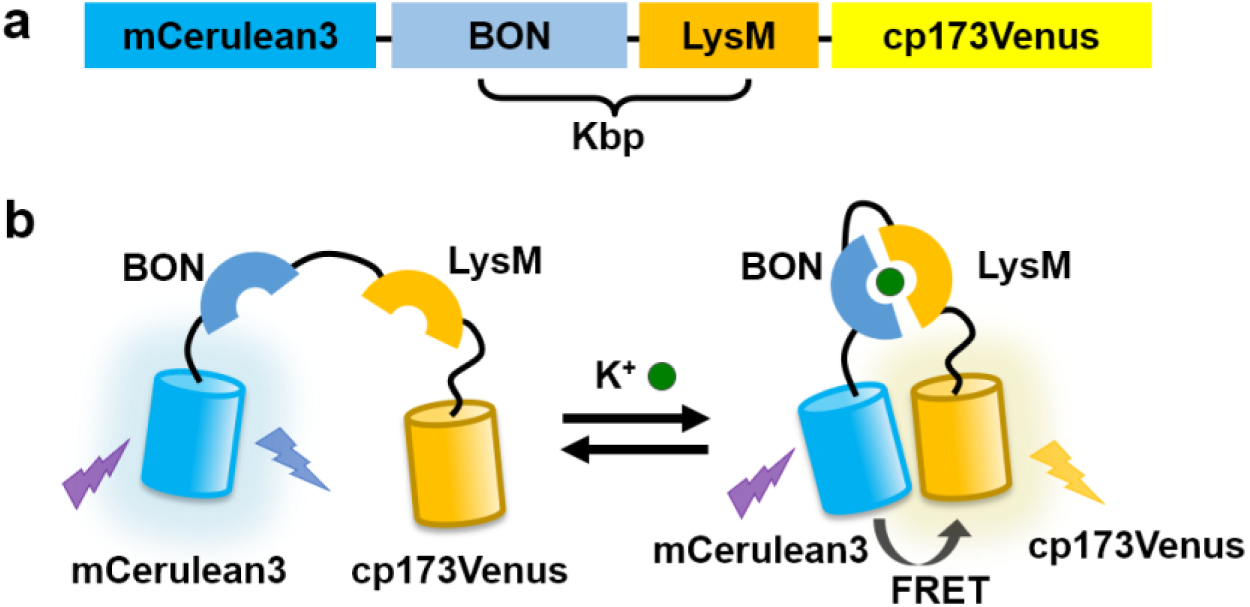
Design of KIRIN1. (**a**) Design and construction of the genetically encoded K^+^ indicator KIRIN1. mCerulean3 (residue 1-228) at the N-terminus is linked with Kbp (residue 2-149), followed by cp173Venus (full length) at the C-terminus. (**b**) Schematic demonstration of the molecular sensing mechanism of KIRIN1. Binding of K^+^ induces a structural change in the Kbp protein, increasing the FRET efficiency between mCerulean3 and cp173Venus in KIRIN1.

### *In vitro* Characterization

To investigate the potential utility of KIRIN1 as a K^+^-indicator, the protein was purified and characterized *in vitro*. To determine the K^+^-induced FRET efficiency change, we first measured the emission spectrum (430 nm to 650 nm) of KIRIN1 protein in Tris buffer (Fig. 2a). Upon the addition of 150 mM K^+^, the mCerulean3 emission (∼475 nm) decreased by ∼50% and cp173Venus (∼530 nm) emission increased by ∼30%, indicating an increase in FRET efficiency resulting from the K^+^-induced structural compaction of Kbp. The maximum FRET acceptor and donor fluorescence emission intensity ratio (F_530_/F_475_) change was 2.3-fold, which is comparable to a number of highly optimized CFP-YFP-based FRET indicators^32,35^.

**Figure 2.**
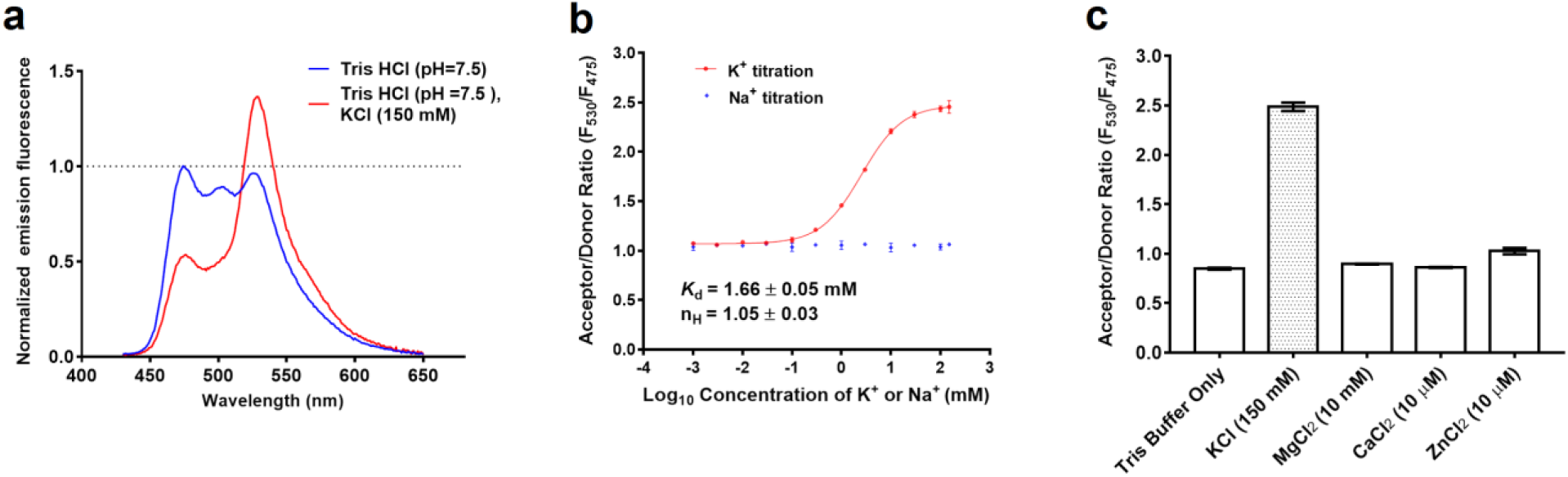
*In vitro* characterization of KIRIN1. (**a**) KIRIN1 emission fluorescence spectrum (excitation 410 nm, emission 430 nm to 650 nm) in Tris HCl buffer with (red) and without (blue) K^+^. (**b**) FRET acceptor-to-donor fluorescence ratio (F_530_/F_475_) of purified KIRIN1 protein at various concentrations (1 μM to 150 mM) of K^+^ (red) and Na^+^ (blue) solutions. (**c**) FRET acceptor-to-donor fluorescence ratio of purified KIRIN1 protein upon addition of various physiologically relevant ions including Mg^2+^ (10 mM), Ca^2+^ (10 μM), and Zn^2+^ (10 μM).

To determine the K^+^ affinity of KIRIN1, we measured the mCerulean3 and cp173Venus fluorescence emission ratio (F_530_/F_475_) over K^+^ concentrations ranging from 1 μM to 150 mM (Fig. 2b). The indicator showed FRET ratio change from 0.1 mM to 100 mM K^+^, which covers the physiologically relevant concentrations. The K^+^ titration curve was fitted with a one-site saturation model, yielding an apparent dissociation constant *K*_d_ = 1.66 ± 0.05 mM and apparent Hill coefficient (n_H_) of 1.05 ± 0.03 (Fig. 2b). To test the selectivity for K^+^ relative to Na^+^, we also incubated KIRIN1 with Na^+^ over the physiological concentration range from 1 μM to 150 mM. The FRET ratio remained unchanged upon the addition of concentrations of Na^+^ up to 150 mM (Fig. 2b), indicating that KIRIN1 has an excellent selectivity for K^+^ over Na^+^. The indicator exhibited no significant FRET ratio change upon the addition of other physiologically relevant ions including Mg^2+^ (10 mM), Ca^2+^ (10 μM), or Zn^2+^ (10 μM) (Fig. 2c), further confirming the specificity of KIRIN1 toward K^+^. In parallel characterization, KIRIN1-GR exhibited an apparent dissociation constant *K*_d_ =

2.56 ± 0.01 mM for K^+^ binding. Like KIRIN1, it also showed a strict selectivity toward K^+^. However, the *in vitro* K^+^ binding–induced FRET ratio change was limited to 20%, much smaller than the 230% ratio change of KIRIN1 (Supplementary Fig.2).

### Imaging of intracellular K^+^ depletion in cell lines

Based on the *in vitro* characterization, Kbp-based genetically encoded K^+^ indicators are promising candidates for visualization of intracellular K^+^ dynamics in live cells. To test the performance of KIRIN1 in cells, immortalized human cervical cancer cells (HeLa) were transfected with the plasmid encoding KIRIN1. We then induced changes in intracellular K^+^ concentrations using pharmacological approaches. Amphotericin B is an antifungal drug that is known to induce K^+^ efflux from cells. Ouabain is an inhibitor of sodium-potassium ion pump (Na/K-ATPase), which is essential to maintain high K^+^ concentrations inside cells. Addition of both amphotericin B and ouabain to cells is expected to deplete intracellular K^+^ content^36,37^. Cells were imaged in both the CFP (excitation 436/20 nm emission 483/32 nm) and FRET (excitation 438/24 nm, emission 544/24 nm) channels, and the FRET ratio image of KIRIN1 was calculated by dividing the FRET channel image by the CFP channel image (Fig. 3a). Upon treatment with 5 μM amphotericin B and 10 μM ouabain, the indicator exhibited a gradual decrease of 15% in FRET ratio over a period of ∼60 minutes, signaling a decrease of K^+^ concentration in the cells (Fig. 3b). The control experiment using KIRIN1-expressing cells treated with DMSO vehicle and imaged under identical conditions revealed no significant change in FRET ratio over the same time period (Fig. 3c). A similar FRET ratio decrease was observed with cells expressing KIRIN1-GR (Supplementary Fig. 3), with a smaller percentage change of 8%. These findings demonstrate that KIRIN1 and KIRIN1-GR are capable of monitoring intracellular K^+^ changes across a population of live cells in real time.

**Figure 3.**
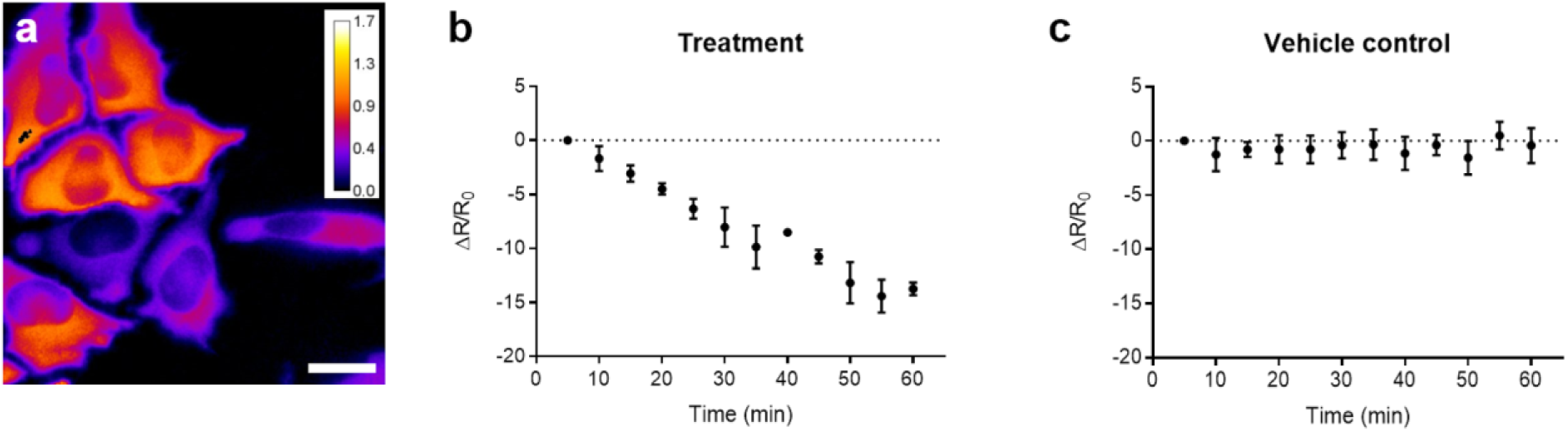
Imaging intracellular K^+^ depletion using KIRIN1. (**a**) Representative FRET acceptor (excitation 438/24 nm, emission 544/24 nm) to donor (excitation 436/20 nm emission 483/32 nm) fluorescence ratio (R = F_acceptor_/F_donor_) image of live HeLa cells expressing KIRIN1 (scale bar = 20 μm). (**b**) Trace of FRET acceptor-to-donor fluorescence percentage ratio change (ΔR/R_0_) after treatment of live HeLa cells expressing KIRIN1with 5 μM amphotericin B and 10 μM Ouabain (n = 7). (**c**) Trace of FRET acceptor-to-donor fluorescence percentage ratio change (ΔR/R_0_) without chemical treatment (n = 13) on live HeLa cells expressing KIRIN1.

### Imaging intracellular K^+^ dynamics in primary neurons

K^+^ plays an essential role in maintaining the proper membrane potential of neurons. The depolarization and repolarization of neurons that underlies neuronal activity are associated with K^+^ efflux/influx, respectively. We next examined whether changes in intracellular K^+^ during neuronal activities can be detected optically using KIRIN1. We utilized primary neurons cultured from the rat hippocampus/cortical region, a well-established neuronal model. KIRIN1 was expressed in neurons via transient transfection, and it diffused evenly throughout the cell and filled neuronal cell bodies and processes (Fig. 4a). Previous studies^38,39^ using electrode-based measurements of extracellular K^+^ concentration revealed that stimulation with glutamate, a major excitatory neurotransmitter, mediates a strong K^+^ efflux from neurons, resulting in depletion of intracellular K^+^. After perfusion of glutamate (1 mM) for a short period (30 s), neurons expressing KIRIN1 underwent a 10% decrease in FRET ratio within 120 s (Fig. 4b). After washing out glutamate, the FRET ratio of KIRIN1 returned to baseline after ∼600 s (Fig. 4b), suggesting the re-equilibration of intracellular K^+^ concentration in neurons.

**Figure 4.**
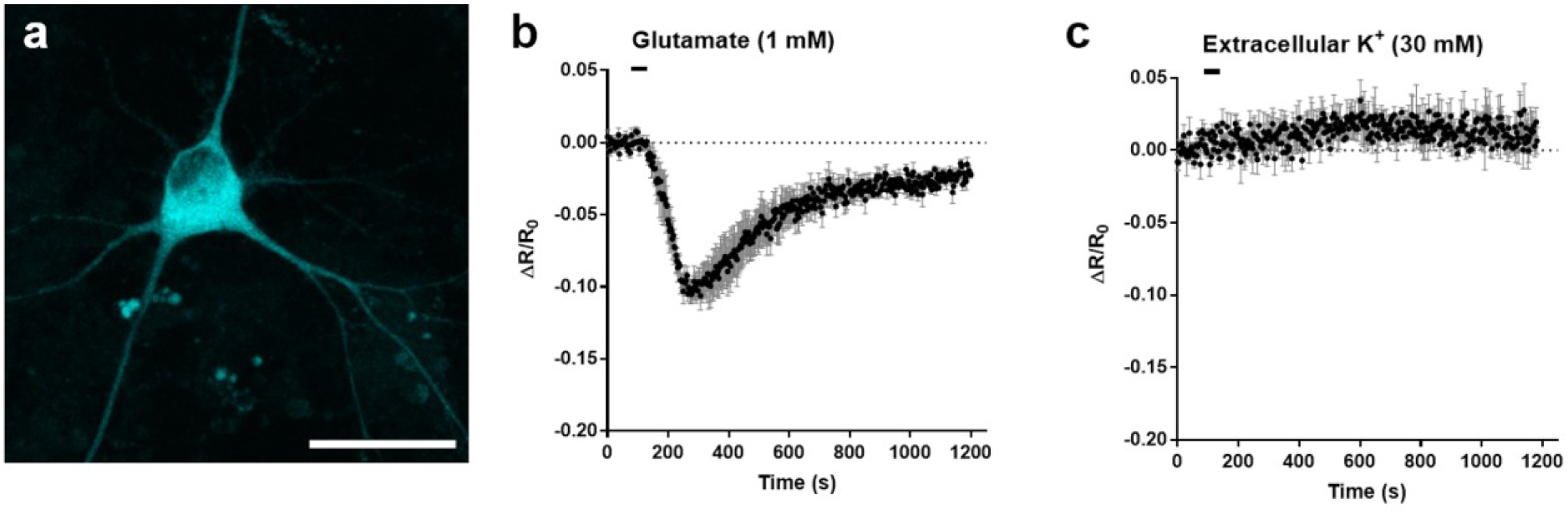
Imaging neuronal K^+^ dynamics using KIRIN1. (**a**) Representative CFP channel fluorescence image of dissociated neuron expressing KIRIN1 (scale bar = 30 μm). (**b**) FRET acceptor-to-donor fluorescence percentage ratio change time course with the treatment of glutamate (n = 6). (**c**) FRET acceptor-to-donor fluorescence percentage ratio change time course with treatment with a high concentration of extracellular K^+^ (n = 3).

To further confirm that the FRET ratio change is due to K^+^ dynamics rather than other cellular changes associated with neuronal activities, we performed the same set of experiments but replaced glutamate stimulation with high extracellular concentrations of K^+^ (30 mM)^40^. This depolarizes neurons but does not significantly change intracellular K^+^ concentrations, possibly due to the fact that the K^+^ gradient across the membrane is reduced and outward K^+^ current is attenuated^38,39^. Stimulating neurons expressing KIRIN1 with high extracellular K^+^ concentration (30 mM) did not result in any detectable FRET ratio changes (Fig. 4c), confirming that KIRIN1 FRET changes specifically reflects K^+^ dynamic during neuronal activity.

## Discussion

Here we designed and developed a genetically encoded K^+^ indicator, KIRIN1, based on bacterial Kbp and the FRET pair of mCerulean3 and cp173Venus. Our characterizations of KIRIN1 demonstrated FRET efficiency changes across physiologically relevant concentrations of K^+^. The dynamic range in the FRET ratio of KIRIN1 *in vitro* is 230%, comparable to some highly optimized FRET-based ion indicators^41–43^. Importantly, KIRIN1 is much more selective for K^+^ than Na^+^, due to the inherent K^+^ selectivity of Kbp^28^. Our findings on KIRIN1 are largely consistent with a recent report of a similarly designed K^+^ indicator, GEPII1.0, which also uses Kbp and the cyan/yellow FRET pair^44^. The main difference between GEPII1.0 and KIRIN1 is the use of mseCFP versus mCerulean3, respectively, as FRET donors. Both indicators use Kbp as a sensing domain and cp173Venus as a FRET acceptor. Together, these studies fully confirm that Kbp-based K^+^ indicators are well-suited for sensing K^+^ *in vitro* and in live cells.

In addition to KIRIN1, we constructed another K^+^ indicator, KIRIN1-GR, based on the green and red FP FRET pair Clover/mRuby2. However, the change in FRET ratio of KIRIN1-GR is substantially smaller than that of KIRIN1, which is like due to the relatively low quantum yield of mRuby2 (0.38)^45^ and the lack of weak dimerization effect of CFP/YFP pair^46,47^. Nonetheless, the GFP/RFP-based, FRET pair–based KIRIN should offer a high signal-to-background ratio with minimal spectral bleed-through. Future efforts to increase the KIRIN1-GR FRET change could be directed at replacing the Clover/mRuby2 FRET pair with newly developed brighter GFPs such as mClover3 or mNeonGreen and brighter RFPs such as mRuby3 or mScarlet^34,48,49^.

KIRIN1’s large FRET ratio change and high specificity make it a promising probe for intracellular K^+^ imaging. Our live cell imaging experiments in cell lines and dissociated neurons demonstrated that KIRIN1 is useful for imaging K^+^ dynamics intracellularly. The advent of Kbp-based K^+^ biosensors would open new avenues for future investigation of cellular signaling resulting from (or causing) normal or abnormal K^+^ dynamics, including neuronal and cardiovascular activity and innate immunity activation, either at the single-cell level or across a large cell population in a high-throughput manner. Expressing such indicators *in vivo* in animal models via viral vectors or through transgenic mice could also offer a new optogenetic tool for understanding the role of K^+^ dynamics *in vivo*.

## Methods

### Plasmid DNA construction

DNA encoding KIRIN1 was synthesized and cloned into the pET28a expression vector by GenScripts. For KIRIN1-GR, the Kbp sequence was synthesized by Integrated DNA Technologies (IDT) as a double-stranded DNA fragment; DNA encoding Clover and mRuby2 was obtained from Addgene (plasmid #49089); the DNA fragments were PCR amplified using Phusion polymerase and then cloned into the pBAD/His-B expression vector (Thermo Fisher Scientific) using Gibson assembly (New England Biolabs). KIRIN1 and KIRIN1-GR genes were PCR amplified, digested, and ligated into the mammalian expression vector pcDNA3.1 (Thermo Fisher Scientific). pcDNA 3.1-NES-KIRIN1 was constructed by PCR with the primer containing the nuclear exclusion signal (NES) encoding the sequence (LALKLAGLDIGS). All constructs were verified by DNA sequencing.

### *In vitro* characterization

The indicator protein was expressed as His-tagged recombinant proteins in *E.coli*. with an induction temperature of 20°C overnight. Bacteria were centrifuged at 10,000 rpm, 4°C for 10 min, lysed using sonication, and then clarified at 14,000 rpm for 30 min. The protein was purified from the supernatant by Ni-NTA affinity chromatography (McLab) according to the manufacturer’s instructions and then buffer exchanged using a PD-10 (GE Healthcare Life Sciences) desalting column. For K^+^ *K*_d_ determination, purified protein was diluted into a series of buffers with K^+^ concentration ranges from 0 to 150 mM. The fluorescence spectrum of the KIRIN1 in each solution (100 μL) was measured using a Tecan Safire2 microplate reader with excitation at 410 nm and emission from 430 nm to 650 nm. The fluorescence spectrum of the KIRIN1-GR in each solution (100 μL) was measured using a Biotek microplate reader with excitation at 470 nm and emission from 490 nm to 700 nm). Titration experiments were performed in triplicate. The FRET ratios (F_Acceptor_/F_Donor_) were plotted as a function of K^+^ concentration. Data was analyzed using Graphpad Prism to obtain apparent *K*_d_ and the apparent Hill coefficient.

### Live cell imaging

For mammalian cell line imaging, HeLa or HEK293 cells were maintained in Dulbecco’s modified Eagle medium (DMEM) supplemented with 10% fetal bovine serum (FBS, Thermo Fisher Scientific), Penicillin-Streptomycin (Thermo Fisher Scientific), and GlutaMAX (Thermo Fisher Scientific) at 37°C with 5% CO_2_. Cells with 60-70% confluence on 35-mm glass-bottom dishes (In Vitro Scientific) were transfected using Polyjet (SignaGen) or Lipofectamine 2000 (Thermo Fisher Scientific) according to the manufacturer’s instructions. Cells were imaged 24 h after transfection. Immediately prior to imaging, medium was replaced with 1 mL of 20 mM HEPES buffered HBSS (HHBSS). Cell imaging was performed with a Zeiss Axiovert 200 microscope with a 40x objective (Figure 3 and Figure 5) or an Olympus IX50 microscope with a 60x objective (Supplementary Figure 3). HeLa or HEK293 cell images were processed using ImageJ.

For imaging of cultured dissociated neurons, dissociated E18 Sprague Dawley hippocampal/cortical neurons were grown on 35-mm glass-bottom dishes (In Vitro Scientific) containing NbActiv4 medium (BrainBits LLC) supplemented with 2% FBS and Penicillin-Streptomycin (Thermo Fisher Scientific). Half of the culture medium was replaced every 4–5 days. Neurons were transfected on day 8 using Lipofectamine 2000 (Thermo Fisher Scientific), according to the manufacturer’s instructions, with the following modifications. Briefly, 1–2 μg of plasmid DNA and 4 μl of Lipofectamine 2000 (Thermo Fisher Scientific) were added to 100 μl of NbActive4 medium to make the transfection medium. Half of the neuron culture medium was removed from each neuron dish and combined with an equal volume of fresh NbActiv4 medium to make a 1:1 mixture and incubated at 37°C and 5% CO_2_. All dishes were replenished with freshly conditioned (at 37°C and 5% CO_2_) NbActiv4 medium. Following the addition of transfection medium, neurons were incubated for 2–3 h. The medium was then replaced using the conditioned 1:1 mixture medium. Cells were imaged 48–72 h after transfection. Fluorescence imaging was performed in HHBSS buffer using a Zeiss microscope with a 40x objective or Olympus FV1000 microscope with a 20x objective (Figure 4). Neuron images were processed using Olympus FluoView.

### Statistical analysis

All data are expressed as mean ± s.e.m. Sample sizes (n) are listed for each experiment.

## Acknowledgements

SYW is supported by a NSERC graduate scholarship and an Alberta Innovates Technology Futures scholarship. Work in the lab of REC is supported by grants from CIHR (FS 154310), NSERC (RGPIN 288338-2010), NIH (U01 NS094246 and U01 NS090565), and Brain Canada. MD acknowledges support from NIH (NS080833 and AI132387), the NIH-funded Harvard Digestive Disease Center (P30DK034854), and Boston Children’s Hospital Intellectual and Developmental Disabilities Research Center (P30HD18655). MD holds the Investigator in the Pathogenesis of Infectious Disease award from the Burroughs Wellcome Fund. We thank the University of Alberta Molecular Biology Services Unit for technical support, Dr. Christopher W. Cairo for providing access to instrumentation, and Dr. Matthew D. Wiens for helpful discussion regarding the manuscript.

## Author contributions

YS performed design, construction, and *in vitro* characterization of KIRIN1 and KIRIN1-GR. YS, SYW, and SIM performed mammalian plasmid construction. YS and SYW performed live imaging in mammalian cell lines. VR and YQ performed imaging in dissociated neurons. YS, KB, REC, and MD supervised the research. YS, REC, and MD wrote the manuscript.

## Data and material availability

The data supporting this research are available from the corresponding authors upon request. Plasmid constructs encoding KIRIN1 and KIRIN1-GR are available through Addgene.

**References**

## Supplementary Figures

**Supplementary Figure 1.**
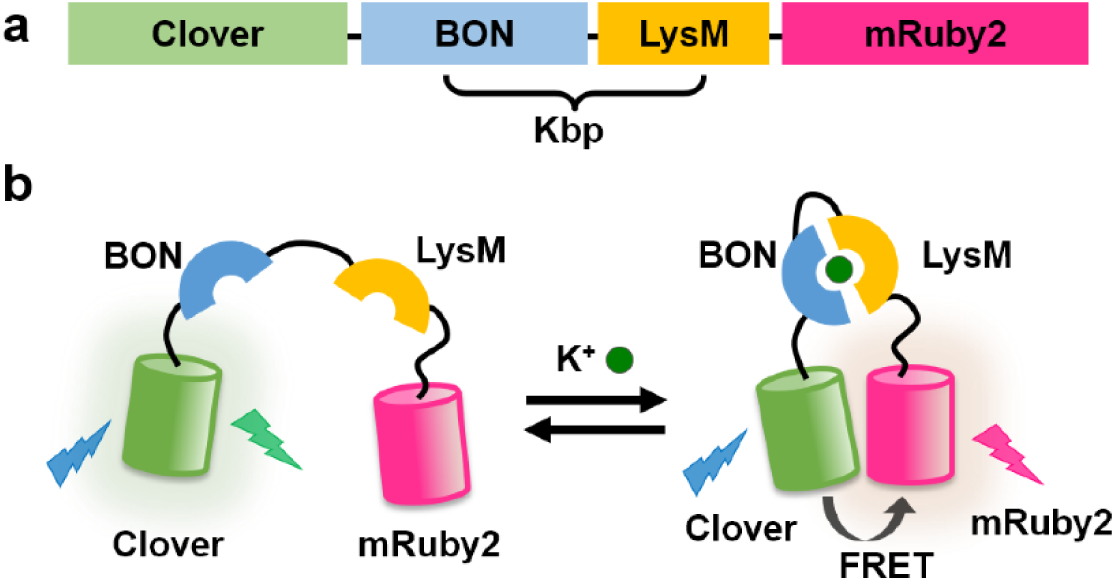
Design of KIRIN1-GR. (**a**) Design and construction of the genetically encoded K^+^ indicator KIRIN1-GR. (**b**) Schematic of the molecular-sensing mechanism of KIRIN1-GR.

**Supplementary Figure 2.**
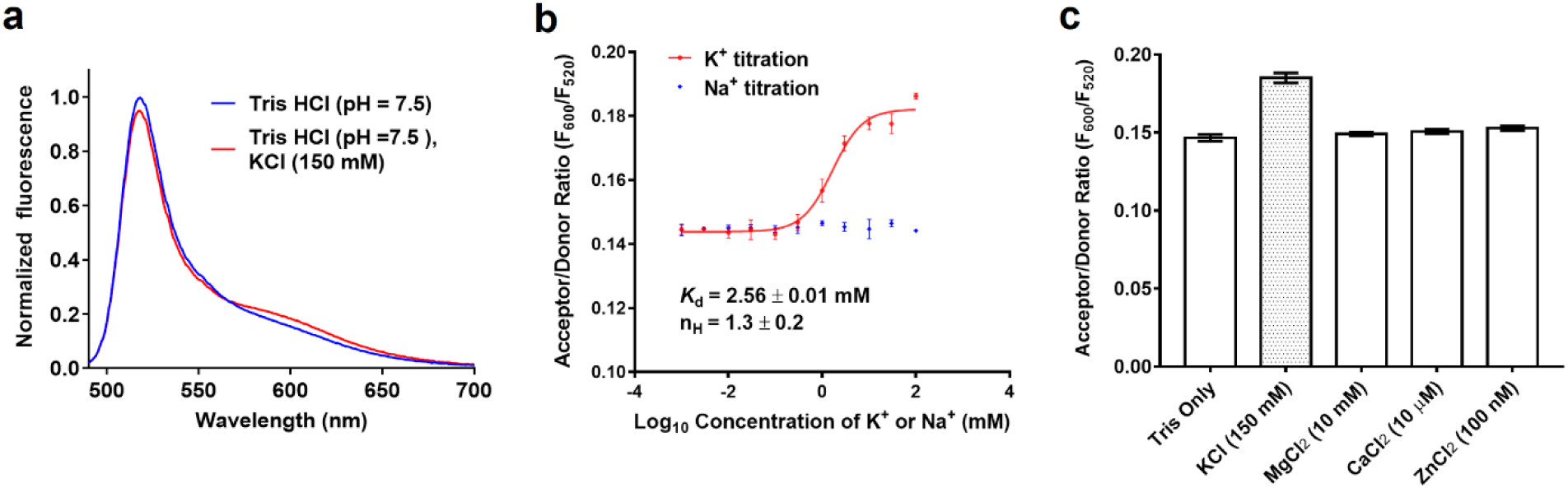
*in vitro* characterization of KIRIN1-GR. (**a**) Emission fluorescence spectrum of KIRIN1-GR with (red) and without (blue) K^+^. (**b**) K^+^ titration curve (red) and Na^+^ titration (blue) of KIRIN1-GR according to FRET acceptor-to-donor fluorescence ratio (F_600_/F_520_). (**c**) FRET acceptor-to-donor fluorescence ratio of the genetically encoded K^+^ indicator elicited by adding different physiologically relevant ions including Mg^2+^ (10 mM), Ca^2+^ (10 μM), and Zn^2+^ (10 nM).

**Supplementary Figure 3.**
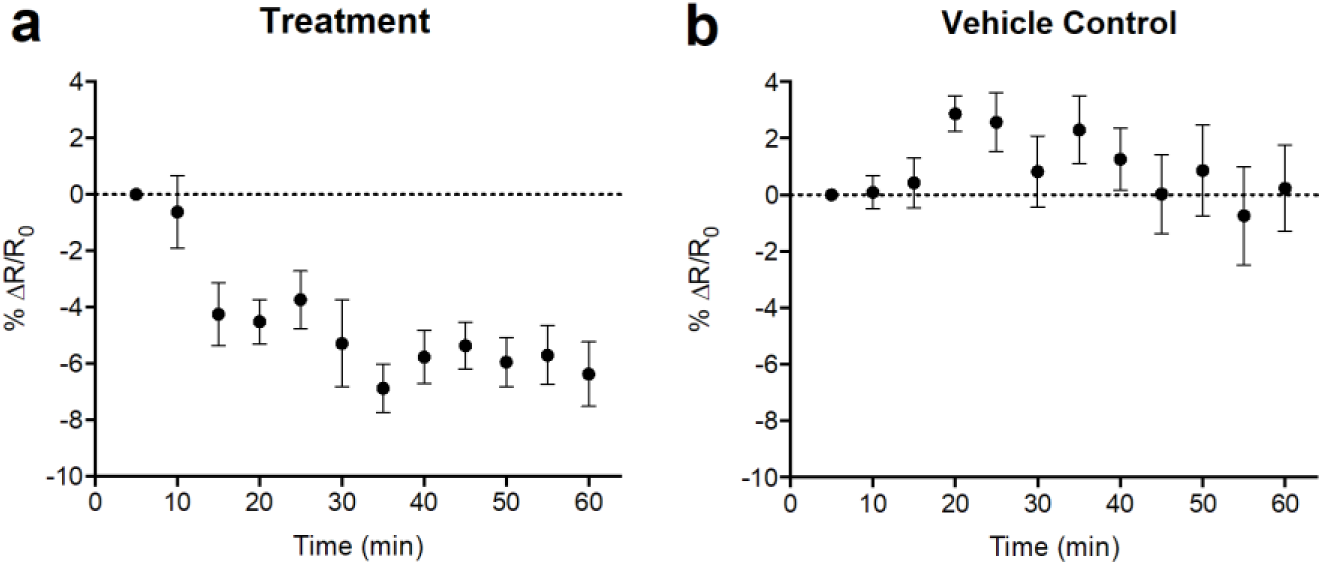
Imaging intracellular K^+^ depletion using KIRIN1-GR. (**a**) FRET acceptor-to-donor fluorescence ratio time course produced by treatment with Amphotericin B and Ouabain (n=5). (**b**) FRET acceptor-to-donor fluorescence ratio time course without chemical treatment (n=7).

